# Mechanistic Insights into the Structural Asymmetry of the LanFEG Transporter NisFEG in Lantibiotic Immunity

**DOI:** 10.1101/2025.10.09.681468

**Authors:** Pablo A. Cea, Julia Gottstein, Stephan Schott-Verdugo, Christian Mammen, Sander H.J. Smits, Holger Gohlke

## Abstract

Nisin is one of the best studied antimicrobial peptides. Still, how nisin-producing strains can protect themselves against nisin’s bactericidal effects is only partially understood. Located within the nisin biosynthesis operon, the heterotetrameric ABC transporter NisFEG transports nisin to the extracellular environment, granting autoimmunity to the producer strain. NisFEG belongs to the LanFEG family of ABC transporters, members of which are found in some lantibiotic-producing bacterial strains. However, their structure has not been elucidated. In this work, we constructed a full atom model of NisFEG in the ATP-bound conformation. The architecture of the complex reveals a narrow contact interface between the two transmembrane chains, with prominent lateral clefts, similar to those observed in other exporters of hydrophobic compounds. Through molecular dynamics (MD) simulations, we observed that one of the most conserved elements of the LanFEG family, the E-loop of the nucleotide-binding domain, interacts preferentially with a small intracellular helix of the NisG transmembrane chain. By combining co-solvent MD simulations and predictions of the binding mode of the terminal segment of nisin, we could identify a putative interaction surface, located predominantly on NisE. Our results suggest that nisin extrusion operates in an asymmetric manner, where contacts between the E-loop and NisG are the driving force for the conformational changes triggered by ATP hydrolysis, whereas the NisE subunit is the main mediator of interactions with the lantibiotic. This functional asymmetry could explain why the LanFEG family has evolved two distinct transmembrane chains, where each one was selected to perform a single step in an optimal way, maximizing the immunity of lantibiotic-producing bacteria.

## Introduction

Bacteria produce antimicrobial peptides (AMPs) that target closely related species for survival. These peptides are produced under stress conditions, including limited nutrient availability or nutrient starvation, and are produced by both Gram-(+) and Gram-(-) bacteria and usually possess a narrow-spectrum antibacterial activity. Bacteriocins are a class of AMPs of particular interest since they can exert different modes of action in both genera.

Lanthipeptides are a class of bacteriocins that are produced by specific biosynthetic clusters mainly in Gram-positive bacteria, but their occurrence is not restricted to this group; they are also found in Gram-negative bacteria [1] and cyanobacteria [2, 3]. They are ribosomally synthesized as pre-peptides and undergo several post-translational modifications (PTMs) and are activated after secretion by leader peptide cleavage via a specific protease. The most distinctive PTMs occur on serine and threonine residues that are dehydrated, resulting in the non-standard α,β-unsaturated amino acids 2,3-didehydroaline (Dha) and 2,3-didehydobutyrine (Dhb) [4, 5], followed by a Michael-type condensation of these amino acids with cysteine residues, yielding a thioether crosslink, so-called lanthionine rings (Lan or MeLan rings) [6, 7].

Within the bacteria, lanthipeptide operons are organized as biosynthetic gene clusters (cluster I-V), in which the precursor peptide (LanA) is located with the enzymes for modification, transport (LanT), processing (LanP), and regulation (LanR and LanK). In class I and II, genes for immunity against lanthipeptides can be found for (LanI and or LanFEG) [8]. Here LanI is encoding a lipoprotein, which binds the lanthipeptide, and an ABC transporter LanFEG, which expels the lanthipeptide from the membrane, inhibiting pore formation [8].

The best described lantibiotic is nisin [9], which is naturally produced by several *Lactococcus lactis* strains. It comprises 34 amino acids, which form five lanthionine rings, and has a broad spectrum of activity, inhibiting the growth of Gram-positive and Gram-negative bacteria (Supplementary Figure 1) [10]. Nisin acts as a membrane-targeting peptide and has two main mechanisms of action: I) it can bind to the pyrophosphate groups of Lipid II, which is an essential carrier for bacterial cell wall synthesis, thereby sequestering synthesis and halting cell division, and II), nisin/Lipid II complexes embedded in the bacterial membrane can oligomerize leading to the formation of lethal pores [11]. Hence, nisin has both bacteriostatic and bacteriolytic effects, which makes it a highly potent (activity in the low nM range) antibacterial compound [12].

Due to its high potency, high stability, and broad target membrane spectrum, nisin has important industrial applications. Currently, it is the only bacteriocin approved by the FDA to be used as a food preservative [13]. Moreover, intensive research has been performed to bolster its clinical use, especially against therapy-resistant bacteria such as methicillin-resistant *Staphylococcus aureus* (MRSA) [14].

In their natural environments, bacteria use antibiotic production as a means to eliminate closely related competition from a given niche [15]. It is therefore essential that they possess adequate molecular machinery to protect themselves from the detrimental effects of their own antimicrobial compounds. For this reason, within the nisin biosynthesis operon, there is a set of genes that are responsible for conferring immunity to the producing strain [16]. These genes encode for a membrane-anchored protein (LanI) known as NisI and an ABC transporter (LanFEG) known as NisFEG [17].

NisI is a 27 kDa protein with a membrane anchoring region located at the N-terminus and two extracellular β-barrel domains [18]. Current evidence points toward this protein acting as a nisin binder, preventing its interaction with lipid II [19, 20]. Structural data suggest that the binding of nisin to NisI occurs through a binding cleft located between the two β-barrel domains [18] but the C-terminal domain alone is also capable of interacting with the lantibiotic [19].

NisFEG is an ABC transporter belonging to the LanFEG family [21]. The complex is formed by three protein chains, encoded by the genes *nisF*, *nisE*, and *nisG*. The functional channel is organized as a heterotetrameric structure with stoichiometry 2(F):1(E):1(G). This arrangement places NisFEG within type V of ABC transporters, according to the classification scheme proposed by C. Thomas *et al.* [22]. NisG and NisE constitute the transmembrane region of the transporter, while NisF is intracellularly located and forms a homodimer. The dimer contains the nucleotide-binding domains (NBD) that hydrolyze ATP to drive the activity of the transporter. Experimental work showed that NisFEG, as well as other members of the LanFEG family, such as NukFEG, actively remove the lantibiotic from the cellular membrane of *L. lactis* [21, 23]. Recently, the nisin-like biosynthetic gene clusters have been identified in a widely diverse set of organisms, including human pathogenic strains [24], where the presence of, e.g., the *lanfeg* genes is used for the identification of these clusters in the genomic background.

To date, no 3D structure has been experimentally resolved for any member of the LanFEG family. Therefore, the mechanistic details of how the transporter works remain unknown. So far, the sequence-based analysis combined with homology modeling has revealed the presence of a conserved E-loop in LanF proteins. This region is a variation of the conserved Q loop motif observed in other ABC transporters and is essential for conferring immunity against nisin, but not for ATP hydrolysis, implying a role in the conformational rearrangements required for effective antibiotic removal [25]. Previous work established that the C-terminus of nisin plays an important role in the recognition between NisFEG and the lantibiotic, as nisin variants lacking the terminal region were still deadly against cells expressing NisFEG [26]. Similarly, nisin variants varying the crucial hinge region in nisin or a natural variant of nisin produced by another strain are less well recognized by NisFEG indicating a certain degree of substrate specificity [27, 28]. This suggested that NisFEG specifically targets the pore formation mode of action, which is mediated by the C-terminus of nisin. However, the lack of structural information on NisFEG, combined with the large size of nisin, has hindered the development of a hypothesis that can explain the function of the transporter.

Here, we constructed a full structural model of the ABC transporter NisFEG in its closed conformation bound to ATP. By using molecular dynamics simulations, we provide first insights into the possible role of the E-loop in the channel’s function. Furthermore, by combining co-solvent MD simulations and mutational studies, we were able to identify a putative binding region for nisin.

## Results

### Architecture of the NisFEG transmembrane domain and full-length model of NisFEG

We constructed a model of the full transporter NisFEG using AlphaFold3. Model quality was assessed through the predicted local distance difference test (pLDDT) and the Ramachandran plot (Supplementary Figure 2). To better interpret the predicted structure, we searched for remotely homologous ABC transporters whose structure-function relationships have been better described, using FoldSeek [31] with NisG as query and the PDB100 [31] as target database. Two bacterial proteins were identified that have transporter functions similar to NisFEG: the exporter of virulent peptides PmtCD from *Staphylococcus aureus* [32] (PDB ID 6XJH) and the O-antigen exporter from *Aquifex aeolicus* [33] (PDB ID 7K2T).

Comparing the generated NisFEG models with the structures of the homologous transporters reveals that the general architecture of the different transmembrane chains (TMB) is conserved. In all cases, the five transmembrane helices within a protein chain have a similar spatial arrangement (Figure 1A). However, the relative orientation of the helices in the oligomeric assembly differs between the two experimentally determined structures. In PDB ID 7K2T, the protein-protein interface is formed by the direct contact of helices αT1 and αT5 of each opposing chain, whereas in PDB ID 6XJH, the contact is mostly mediated by the two αT5 helices of each chain (Figure 1B). A cross-section, perpendicular to the helices at the center of the membrane, reveals a significantly smaller contact surface area between the protein chains in PDB ID 6XJH than in PDB ID 7K2T (1193.9 Å^2^ vs. 1384.7 Å^2^, respectively, assessed by PDBePISA [34]) (Figure 1C). Furthermore, this difference in helical arrangement leads to the presence of a large cleft located between the chains of PDB ID 6XJH. The model of the NisE:NisG assembly has a similar transmembrane arrangement to 6XJH, with αT5 and αT1 farther apart from each other, and an interface dominated by the interaction of αT5 of the opposing chains, leading to pronounced lateral clefts. Despite the low sequence identity between NisG and NisE (19.67%, aligned with ClustalW), they are structurally very similar and are arranged symmetrically with respect to one another.

**Figure 1.**
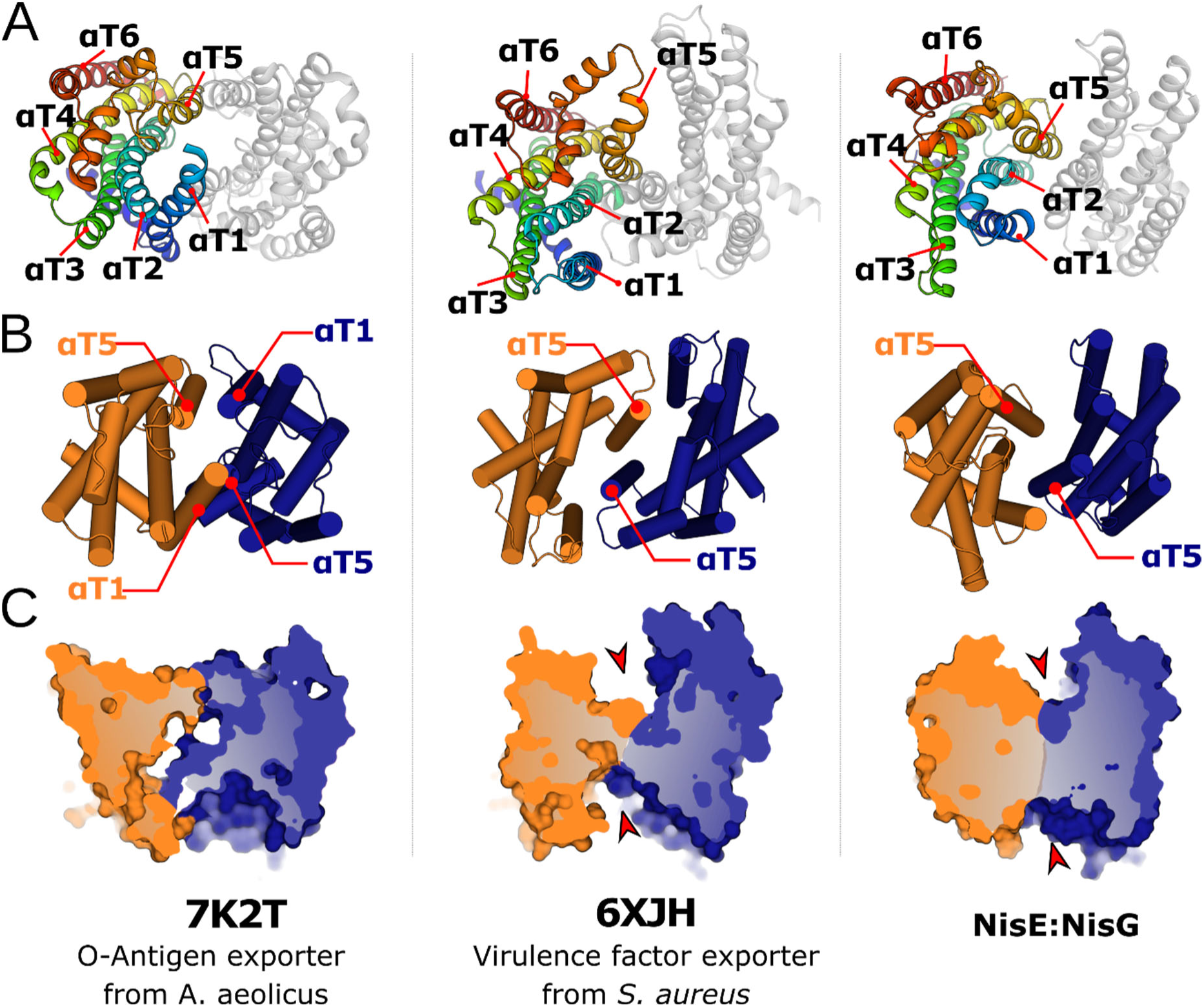
Architecture of the transmembrane region of NisFEG. A) Experimentally determined structures of the the O-antigen transporter Wzm/Wzt from *Aquifex aeolicus* (left; PDB ID 7K2T) and the virulence factor exporter PmtCD from *Staphylococcus aureus* (center; PDB ID 6XJH), and the structural model of NisFEG. For each model, the left chain (identical chains in the case of PmtCD and Wzt, NisE shown for NisFEG) is colored from the N-terminus (blue) to the C-terminus (red) and membrane-spanning helices are labeled from αT1 to αT5. B) For each model, a cylindrical cartoon representation of the transmembrane region is shown highlighting the interfacial helixes. In the case of NisFEG, NisE is colored orange, while NisG is colored blue. C) Perpendicular cross-section of each complex. Pronounced clefts are highlighted with red arrowheads. The same coloring scheme applies as in B).

### Assessing the functional relevance of the conserved E-loop

The LanFEG family possesses a unique structural element known as the E-loop. The name of the loop derives from a conserved Glu residue at its center. So far, this residue is the only amino acid that has been experimentally shown to be essential for the function of the tetramer, but not for ATP hydrolysis [25]. To investigate the structural role of this residue, we performed microsecond-long all-atom molecular dynamics (MD) simulations of the full NisFEG model with ATP-Mg bound in the NBD domains and embedded in a DMPC membrane bilayer. This membrane composition has been successfully used in experimental *in vitro* characterizations of other ABC transporters from *L. lactis* [36]. The simulations reach a state of low-structural variability after ∼200 ns (Supplementary Figure 3).

The E-loop is located where the TMB and the NisF subunits meet (Figure 2A). The loop sits close to small intracellular helical segments present in both TMB chains. Here, Glu76 in one of the NisF subunits can form electrostatic interactions with the residues His17, Asn72, and Gln75 located in a short intracellular helix of NisG (hence we will refer to this subunit as NisF_G_, and the opposite subunit as NisF_E_), whereas on the other subunit (NisF_E_), only the residue Gln76 is oriented in a way that could potentially interact with the E-loop residue (Figure 2B). As expected, the structural model of the complex shows that the E-loop residue Glu76 does not interact directly with ATP in the nucleotide-bound conformation (Supplementary Figure 4), consistent with the NisF homolog NukF retaining its catalytic activity when the corresponding residue is mutated [25].

**Figure 2.**
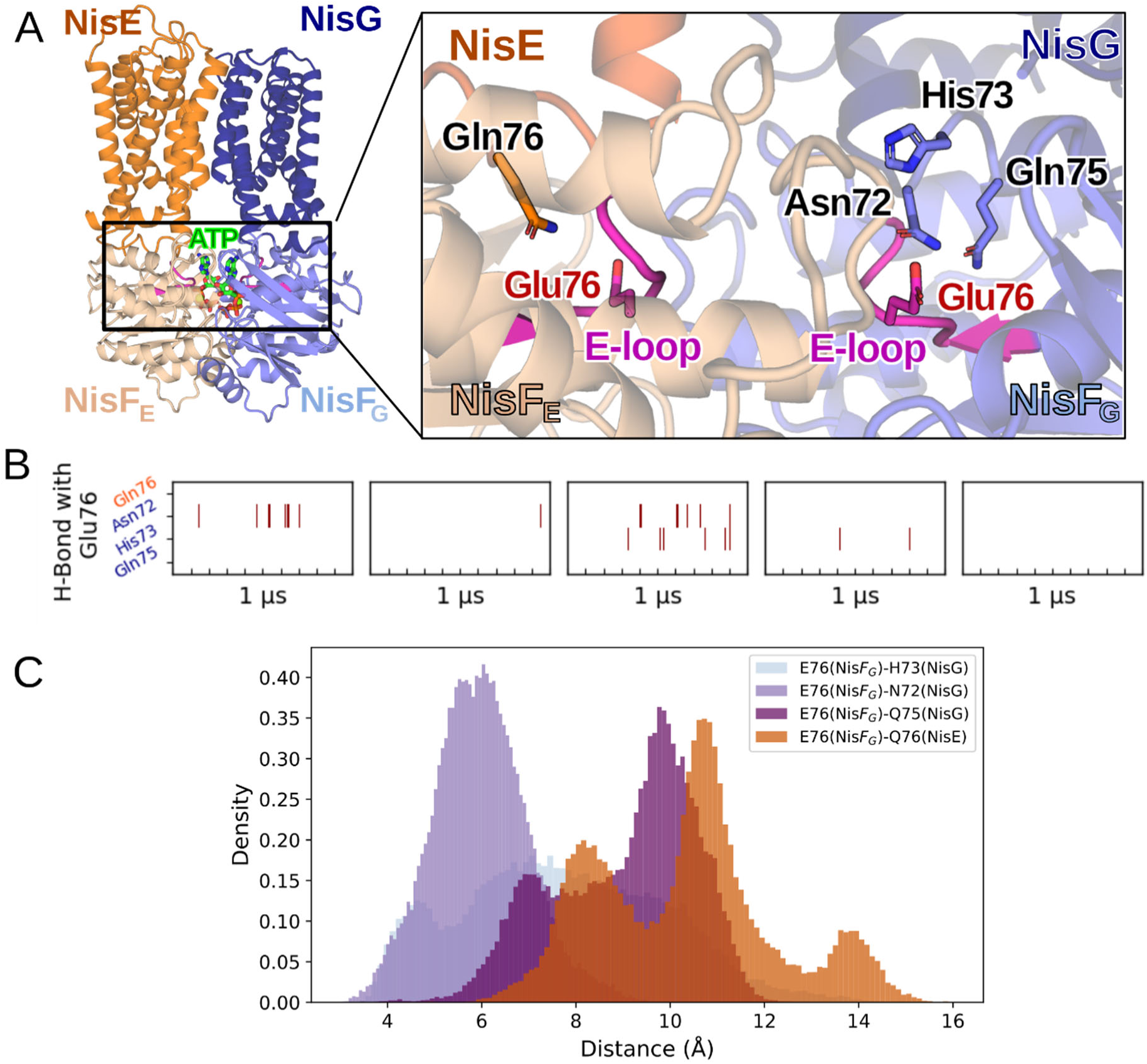
The E-loop mediates interactions with the transmembrane chains of NisFEG. A) Global view of the NisFEG complex model, with the E-loop of both NisF chains highlighted in pink and the ATP molecules shown as green sticks. The right panel shows a close-up of the E-loop. The surrounding residues are shown as sticks and labeled accordingly. B) Time-series showing the presence of hydrogen bonds between Glu76 of the E-loop in NisF_G_ (marked in blue at the Y-axis) and NisF_E_ (marked in orange at the Y-axis). The red stripes represent when the residue pairs are forming an effective hydrogen bond (distance < 3.5 Å between hydrogen and the acceptor and angle > 135° for acceptor-hydrogen-donor) across each replica. C) Histograms showing the pairwise distance distribution between Glu76 and its possible interacting partners across the aggregated simulation time; color code as in panel B.

To evaluate the persistence of interactions mediated by the E-loop residues Glu76, we assessed the presence of hydrogen bonds (defined by a distance cut-off of 3.5 Å and an angle cut-off of 135°) between each Glu76 residue and its adjacent TMB residues (Figure 2B, C). The residue Gln76 from NisE does not establish hydrogen bonds with Glu76 NisF_E_ in any of the replicas. On the other hand, we do observe sporadic, albeit sparse, H-bond formation between Glu76 NisF_G_ with Asn72 and His73, but none with Gln75. We next calculated the distances between the carboxylate group of Glu76 and the functional groups of the interacting residues to determine whether they fall within a range that could still provide electrostatic interactions. From the observed residues, Asn72 stays the closest to Glu76 NisF_G_, with approximately half of the sampled time spent at distances below 6 Å. It is followed by His73, which has a broad distribution, with significant time spent below 6 Å, but reaching also larger separations. Gln75 shows the highest separation to Glu76 from all of the NisG residues, being only surpassed by Gln76 NisE, which spends no time at distances below 6 Å from its possible interaction partner.

To further support the functional relevance of the intermolecular interactions formed by Glu76 and the TMB chains, we analyzed the degree of evolutionary conservation of the involved residues. As the E-loop has already been established as a conserved element within LanFEG transporters, we focused on the interaction partners located at the small helical segments of NisG and NisE. To do this, we employed GEMME [37], a program that predicts mutational outcomes by modeling the evolutionary history of sequences through the construction of phylogenetic trees. We hypothesize that if the observed interactions between the intracellular helix of the TMBs and NisF are relevant, they should exhibit greater conservation, and therefore, the GEMME score should reflect this by a higher probability of functional disruption upon mutation. Residues Ala72 and Phe75 are the most conserved on NisE. Both residues point away from NisF_E_, acting more as hydrophobic contact points to the membrane, while Gln76 has only moderately low scores. On the other hand, Asn72 and Gln75 on NisG display low GEMME scores (Figure 3A), indicating that these sites are likely to disrupt protein function if mutated, correlating with their proximity towards Glu76 NisF_G_.

**Figure 3.**
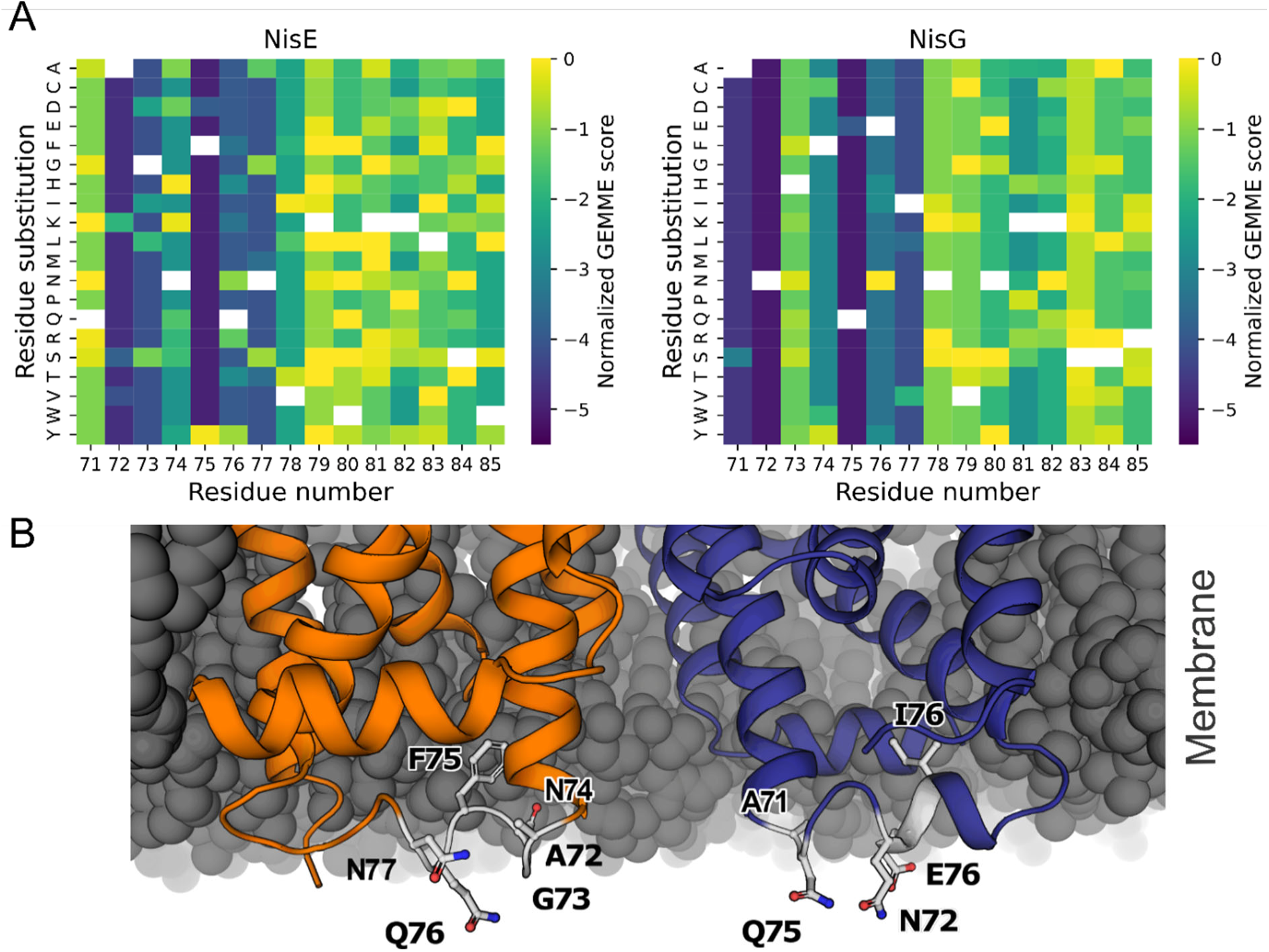
Solvent-exposed residues on the intracellular helix of NisG are more conserved, and their loss could lead to functional disruption. A) GEMME score for residues belonging to NisE intracellular helix. Lower scores indicate a higher degree of conservation and, therefore, a higher likelihood of a residue being functionally relevant. B) Cross-section view of the inner leaflet of the membrane, showing the intracellular section of NisE (orange) and NisG (blue). Residues with low GEMME scores belonging to the intracellular helices are highlighted as sticks -

**Figure 4.**
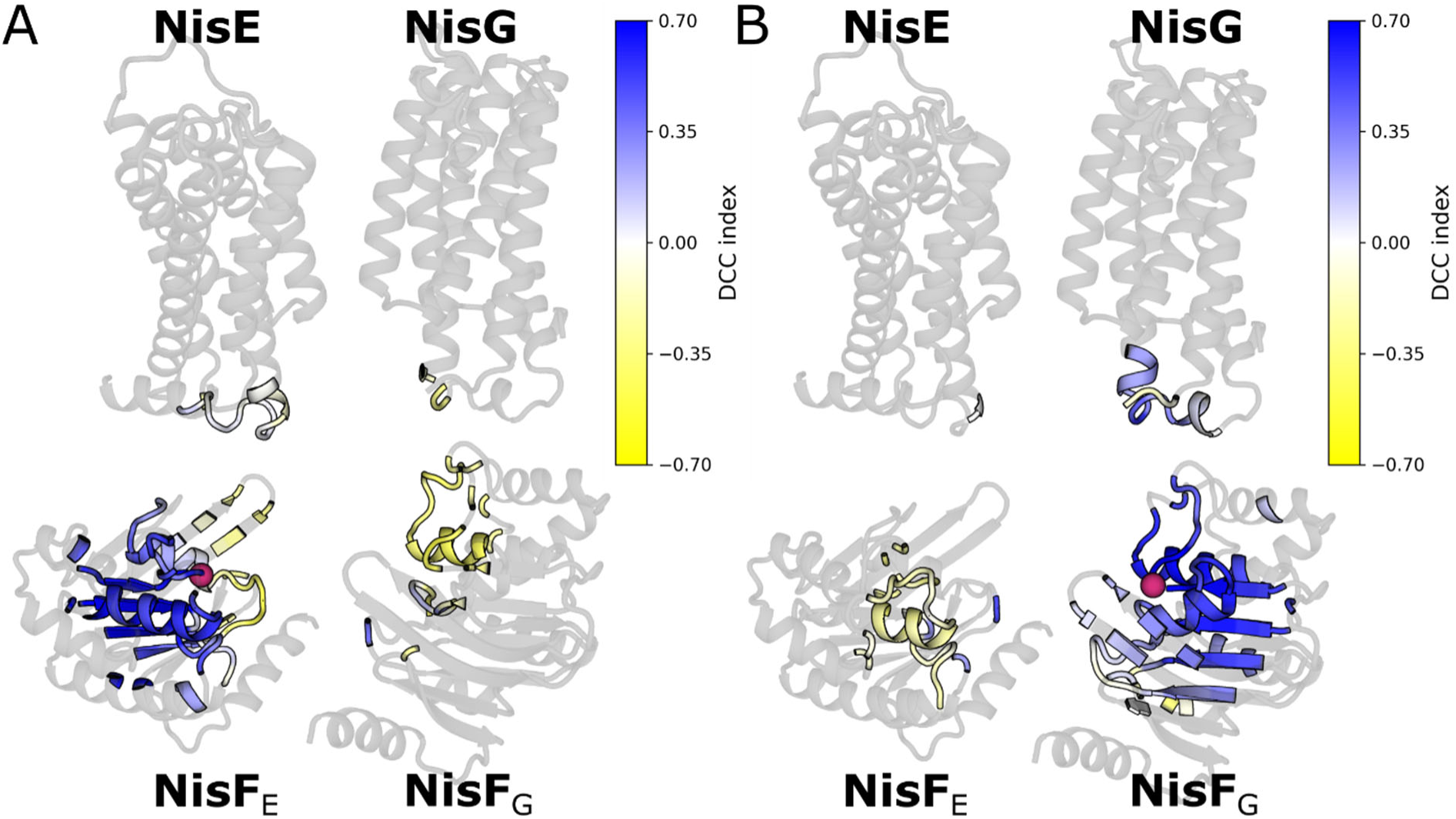
Movements of the E-loop are correlated to those in NisG but not in NisE. A) View of each individual chain of NisFEG. Residues within a radius of 15 Å of the central E-loop residue Glu76 (C_α_ atom highlighted as orange sphere) are colored according to their dynamic cross correlation (DCC) index. The dark blue regions are those moving colinearly with Glu76. B) Same as in A, but highlighting the residues surrounding Glu76 in the E-loop of the second NisF subunit. The same color scale applies to both panels.

To experimentally corroborate our findings, we mutated the interacting partners of the E-loop (Asn72 and Gln75 in NisG) to alanine. As a control, we also mutated the most prominent and conserved polar residue of the equivalent segment of NisE, Gln76, for which GEMME also predicted a high likelihood of functional alteration upon mutation. The complex variants were expressed in *L. lactis*, and the survivability against nisin was measured. Replacing Gln76 with alanine in NisE does not affect the functioning of NisFEG (Table 1). However, the mutation NisG_N72A_ and NisG_Q75A_ led to a reduction of the base activity of about 25% and 17 %, respectively.

**Table 1.** Site-directed mutagenesis at the intracellular NisG helix impairs NisFEG function.

|  | NisFEG<br>control | NisF<sub>H118A</sub> | NisF<sub>K129A</sub> | NisG<sub>N72A</sub> | NisG<sub>Q75A</sub> | NisE<sub>Q76A</sub> |
| --- | --- | --- | --- | --- | --- | --- |
| IC<sub>50</sub><br>(nM) | 118 | 12 | 76 | 93 | 98 | 118 |
| F. o. R<sup>1</sup> | 10 | 1 | 6 | 8 | 8 | 10 |
| Activity<br>(%) | 100 | 0 | 64 | 76 | 83 | 100 |

<sup>1</sup>. Factor of resistance.

To further explore what could be the structural implications of the E-loop-mediated interactions for the system, we calculated a dynamical cross-correlation matrix (DCCM) for the C_α_ atoms of the whole complex. The DCCM allows us to identify which elements within a protein are moving in a synchronous (correlated) way throughout an MD trajectory [38]. A cross-correlation index equal to 1 indicates that the movement of the atom pair is correlated, an index of -1 indicates that the movement is anti-correlated, and an index of 0 indicates that the movements of the atom pair are independent from one another. Therefore, interactions that are strong enough will cause correlated movements of the interacting elements. Figure 3 shows the cross-correlation indexes for all residues within a radius of 15 Å of Glu76 of each subunit, mapped onto each chain of the complex. The strongest correlations for the first NisF subunit are observed in the short intracellular helix of NisG, where residues Asn72 and Gln75 are located. On the other hand, the correlation index between Glu76 of NisF_E_ and its equivalent helical segment in NisE is much lower, and stronger correlations occur only within the NBD itself, which further supports that there is no strong interaction between the E-loop in the second NBD and NisE.

Additionally, we tested whether the presence of ATP-Mg influences the observed behaviour of the E-loop. For this reason, we constructed an alternative model in the apo state and performed equivalent MD simulations. The model showed slightly lower pLDDT values for the residues at the interface between the transmembrane regions and the NBDs (Supplementary Figure 2), and more structural variation was observed in the latter during the MD simulations (Supplementary Figure 5). Both models show an almost identical conformation (RMSD_backbone_: 0.6 Å, Supplementary Figure 6A). The contacts mediated by the E-loop are also similar, with only a deviation of the rotamer for Asn72 NisG. The apo system shows a similar interaction pattern as the ATP-bound state, with more H-bonds between NisG and NisF_G_ observed than between NisE and NisF_E_ (Supplementary Figure 6B). However, in two of the MD replicas, sporadic contacts to Gln76 from NisE occur, while Asn72 NisG does not interact directly with the E-loop. Overall, only slight deviations between the two models are observed.

Lastly, we checked whether the E-loop could also establish contacts relevant for oligomerization between the NBDs. Here, we identified Lys129 on NisF with the potential to form an electrostatic interaction with Glu76 of the opposing NisF subunit (Supplementary Figure 7A). The interaction is very dynamic, with the distance between both residues oscillating between 4 and 10 Å, without a noticeable asymmetry between chains. To assess its importance, we mutated Lys129 to alanine, which results in an activity reduction of 36% (Table 1).

Together, these results show that the E-loop is an asymmetric interface element that directly interacts with NisG but does not display the same interaction pattern with NisE.

### Co-solvent MD simulations to identify putative regions in NisFEG interacting with nisin

So far, there is no evidence on where and how NisFEG binds to nisin. To generate a hypothesis of where the binding interface might be located, we employed cosolvent MD simulations [39]. In this technique, small molecule probes that mimic the main chemical moieties of a ligand are placed in the simulation box and allowed to freely diffuse and interact with the protein surface. Since ligand binding sites provide a chemical environment favorable for non-covalent interactions, the small probes will accumulate in them. This technique has been proven effective to identify previously uncharacterized orthosteric and allosteric sites [39, 40]. Since nisin is a highly hydrophobic molecule and rich in peptide bonds, we utilized benzene, *N*-methylacetamide, acetone, and methanol as probes (Figure 5A). We performed five independent simulation replicas of 1 µs length per probe in the same explicit membrane used above, and then analyzed in which regions of the complex the accumulation of the different probes overlapped. The convergence of the simulation results was assessed by monitoring the radial distribution function of each probe, setting a mask for all the heavy atoms of the probes and the backbone atoms of the protein, and by inspecting the change of normalized density pattern over the simulation course (Supplementary Figure 8).

**Figure 5.**
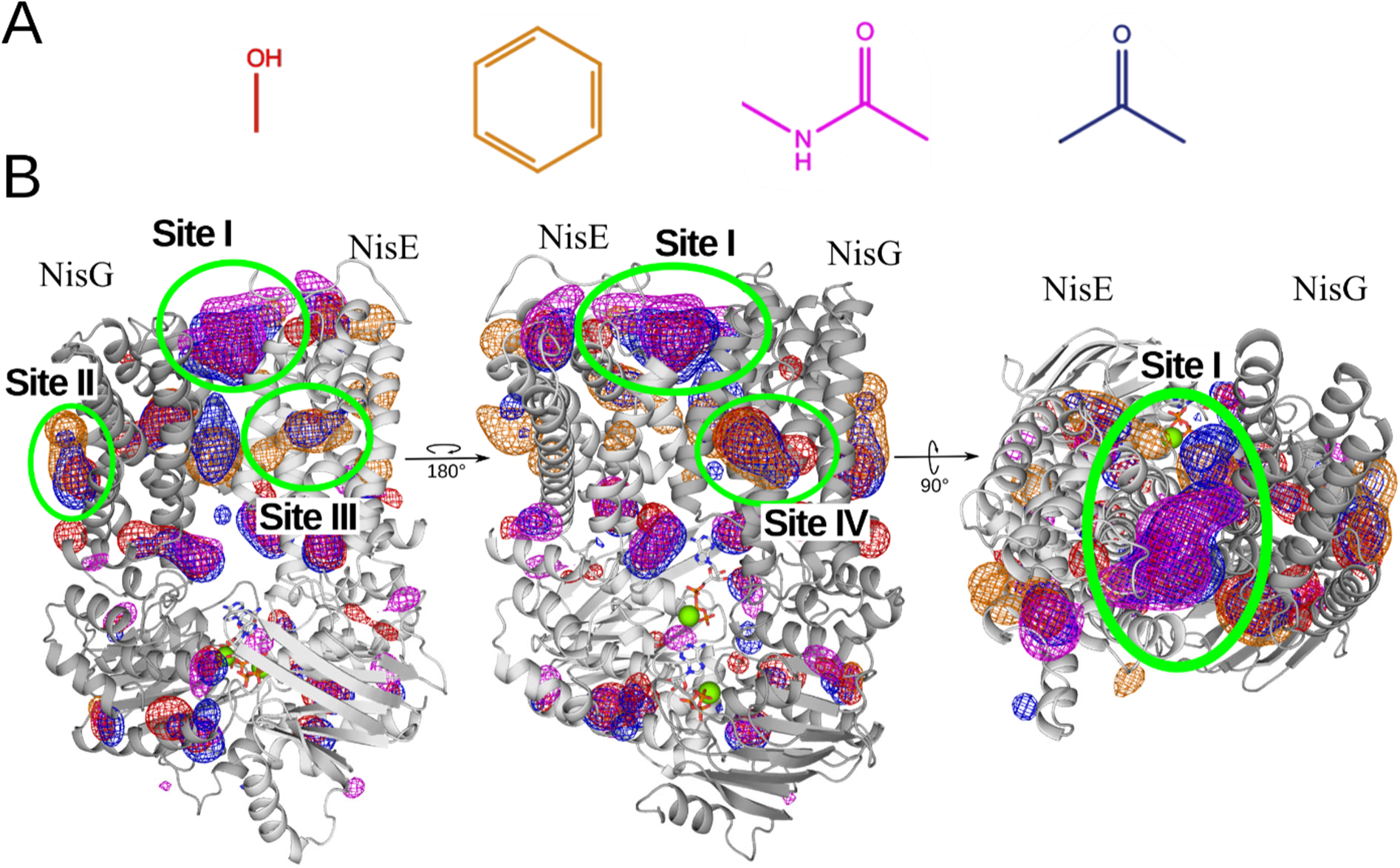
Cosolvent MD simulations reveal putative nisin interaction sites. A) Structure of the used cosolvent probes. Red: Methanol. Orange: Benzene. Magenta: *N*-Methylacetamide Blue: Acetone. B) Density map of the cosolvent probes across the NisFEG complex. Larger densities are observed within the transmembrane region of the transporter. Regions where three or more probes accumulate are highlighted with a green circle and labeled. There are four main sites: one located at the extracellular side of the transporter, at the interface between NisE and NisG (site I), another at the periphery of NisG (site II), and two sites in the transmembrane clefts, one in NisE (site III) and one in NisG (site IV).

Overall, NisFEG accumulates more probes at its transmembrane region than at its intracellular domains. We could identify four regions where there is a marked simultaneous colocalization of at least three probes. The first and largest one (Site I) is located at the extracellular face of the transporter, right at the cleft between NisE and NisG. This site shows a high affinity for the backbone mimetic probes, with minor densities for both the polar and hydrophobic probes. The second site (Site II) is smaller, located on the external side of NisG. The last two sites (Site III and IV) are situated within the lateral clefts of the transporter, one at each side. Site III is slightly smaller and located within the helices of NisE. Interestingly, between this site and Site I, we observe a small accumulation of acetone and benzene. Site IV is larger, with a broader affinity for methanol. These results show that the transmembrane surface of NisFEG has sites with the right physicochemical properties that could possibly form interactions with a peptidic ligand.

### Modeling the interaction between NisFEG and the C-terminus of nisin

Previous work has demonstrated that upon removal of the C-terminus of nisin (C-Nis), NisFEG can not protect the cell against it, implying a direct interaction between the transporter and this part of the lantibiotic [26]. Given that (C-Nis) comprises only six residues (Supplementary Figure 1), current ML-based predictors for biomolecular interactions could infer a possible binding mode. We therefore used Boltz-2 to predict the structure of the full NisFEG complex in the presence of C-Nis. To ensure a proper sampling of the conformational solution space, we generated 600 structural models of the full system.

To infer a putative binding mode, we first clustered the generated poses to identify whether there are clear trends within the data. Since C-Nis is a peptide with a high number of degrees of freedom, clustering only by ligand RMSD can result in significant information loss; thus, we used the coordinates of each ligand’s atom and performed a dimensionality reduction through t-distributed Stochastic Neighbor Embedding (t-SNE). t-SNE generates a nonlinear embedding optimized to preserve local neighborhood relationships of multidimensional data. The distribution of each possible C-Nis configuration on the t-SNE projection space is shown in Figure 6A. There, data points were additionally clustered by their proximity using a Gaussian mixture model and colored accordingly. All the sampled solutions converge to the same location of the NisFEG complex, which is on the extracellular face of the cleft formed by the interface of the transmembrane subunits NisE:NisG. The data is divided into two large clusters, i) and ii), which represent the two possible N- to C-terminus orientations that the lantibiotic could adopt (Figure 6B), of which group ii) occurs 2.8 times more frequently. From this main orientation, we identified the three largest subclusters with 84, 71, and 68 poses, respectively. The subclusters are cohesive groups of poses with average pairwise RMSD values below 3.3 Å each.

**Figure 6.**
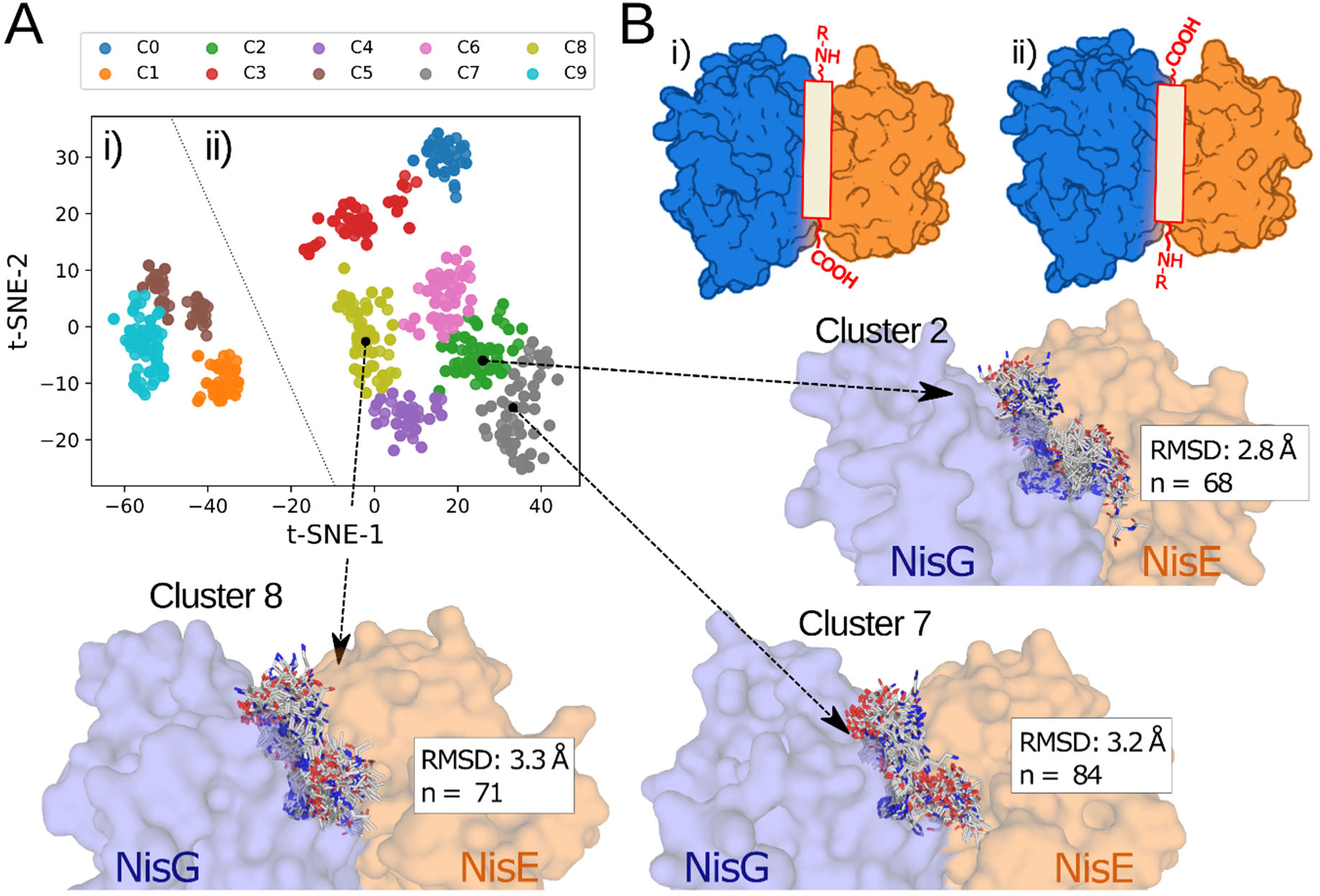
Boltz-2 sampling of C-Nis shows a putative binding site on the extracellular cleft between NisE:NisG. A) t-distributed Stochastic Neighbor Embedding projection derived from the coordinates of the C-Nis atoms when co-modelled with NisFEG. The data is split into two large clusters, labeled i) and ii). The datapoints are colored according to subclusters identified by GMM. Dashed arrow lines point to the ensemble of structures belonging to the three largest sub-clusters. B) Schematic representation of the two possible orientations of C-Nis corresponding to each one of the main clusters (i) and ii) in panel A.

From these subclusters, we selected the pose corresponding to the medoid of the projections as representative to analyze in detail the binding mode, showing the interacting amino acid residues within 5 Å of C-Nis (Figure 7). In all of the observed poses, we observe a larger proportion of residues of NisE surrounding the ligand than of NisG. Furthermore, the position of the His residue of C-Nis is conserved in the three clusters, being flanked by the aromatic residues Phe172_NisE_, W48 _NisE_, and Y47_NisE_. Also of note, in all of the clusters, the N-terminus of the peptide, which would represent where the rest of the nisin molecule would be attached, is placed in an exposed position at the edge of the NisE:NisG interface, implying that the rest of the molecule might fit onto one of the lateral clefts of the transporter. Coincidentally, in poses from group ii), the N-terminus position would leave the rest of the nisin molecule pointing towards where the previously observed Site III on NisE was observed.

**Figure 7.**
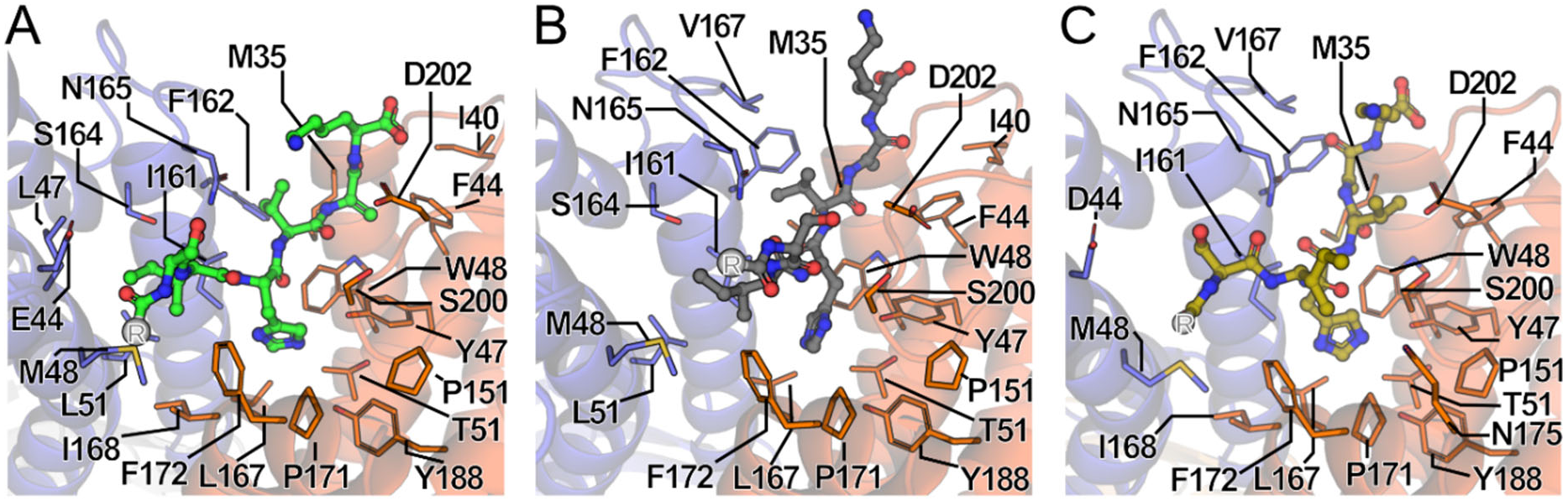
Binding mode of C-Nis derived from the representative poses of the main clusters. Residues within 5 Å of C-Nis in NisG (blue) and NisE (orange) for each one of the largest sub-clusters shown in Figure 6A. The cap atom is highlighted as a gray sphere labeled “R”. A) Subcluster 2, C-Nis shown as green sticks. B) Subcluster 7, C-Nis shown as gray sticks. C) Subcluster 8, C-Nis shown as olive sticks.

## Discussion

In this work, we constructed a full model of the lantibiotic transporter NisFEG from *L. lactis* and focused our analysis on two unknown aspects of it: what is the role of the conserved E-loop, and where could the putative binding site of nisin be located.

A structure-based search using NisG as the query allowed us to identify a distant homolog that also exports a highly lipophilic membrane-disrupting peptide, the ABC transporter PmtCD; a sequence-based search would not have allowed us to identify the homolog due to a sequence identity of only 15.5%. In both cases, the architecture of the transmembrane region of the complex follows a similar arrangement, resulting in a narrow interface with prominent clefts at the sides. Zeytuni *et al.* highlighted the possibility that these side clefts could serve as binding sites for the target peptide [32]. This is a reasonable hypothesis given that the substrate of PmtCD is a hydrophobic helical peptide that embeds itself within the lipid bilayer. This makes the conventional inward-open cavity commonly observed in other ABC transporters for soluble substrates unnecessary, since the peptide cargo already resides in the membrane. Furthermore, eukaryotic ABC transporters that export large hydrophobic substances, such as the ABCA1 transporter, also display a narrow interface mediated mostly by a single helix and large lateral surface clefts [41]. Moreover, the cryo-EM structure of this distantly homologous transporter revealed lipid-like densities at the lateral clefts. It has also been proposed that other ABC transporters, including ABCB4, can function as phospholipid floppases through interactions mediated by the lateral helices [42]. Overall, the current knowledge on ABC transporters suggests that whenever hydrophobic substrates need to be translocated, they can be transported directly from the membrane through interactions with the side of the protein. Thus, it is likely that the whole LanFEG family might follow a similar mechanism.

One aspect that makes transporters of the LanFEG family different from other members of the type V class of ABC transporters is the presence of two distinct transmembrane chains instead of a homodimeric transmembrane region. NisG and NisE share a low sequence identity of 20%, but the heterodimer is required for a functional transporter, suggesting that natural selection has led to the divergence of the two chains. Interestingly, our MD simulations revealed that one of the most conserved elements of the LanFEG family, the E-loop of the NBD, interacts differentially with either transmembrane subunit, even though there are two copies of the loop at equivalent positions within the complex. The E-loop is located near a short intracellular helix present in both transmembrane subunits, but only NisG forms direct polar contacts with the central Glu76 of the E-loop. Furthermore, the helical residues of NisG are more conserved than the equivalent positions of NisE. These conserved interactions apparently have a structural effect: the coupling of motions between the E-loop of NisF_G_ and NisG. Given that E76 is essential for the channel function, the E-loop element likely plays a key role in the propagation of conformational rearrangements triggered by the hydrolysis of ATP towards the transmembrane region that are required for effective nisin removal, thus playing a similar role as the Q-loop in other ABC transporters [43]. Furthermore, experimental work on the human multidrug resistance transporter ABCB1 has shown functional redundancy in the Q-loop; if only one chain is mutated, the transporter retains its activity [44]. This means that it would be feasible for LanFEG transporters to function relying on a single half of the complex to power the required conformational rearrangements. However, the precise nature and extent of conformational rearrangements occurring at the transmembrane region remain open questions within the LanFEG family.

Furthermore, we also explored the influence that ATP binding might exert on the structure of NisFEG. In other ABC transporters, the equivalent loop, known as the Q-loop, has also been proposed to play a role in nucleotide sensing by interacting with the γ-phosphate [45]. Modelling of NisFEG in the apo and ATP-bound states revealed no significant reorientations of the E-loop; however, it is critical to point out that deep learning-based structure prediction methods are limited in predicting the conformational diversity of mobile proteins [46]. Even though we see some slight changes in hydrogen bond patterns between the TMB and the E-loop in the apo model, the residues involved also have a lower pLDDT score, and therefore lower confidence in the accuracy of the model, meaning that the differences could also just be a byproduct of modelling inaccuracies and not a real effect caused by the ATP.

The lack of structural information for LanFEG transporters, combined with the large size and complexity of nisin, has hindered the proposal of a putative binding mode. Our first step to uncover a possible nisin-binding region within NisFEG was to conduct co-solvent MD simulations with probes that mimic the main features of nisin, an approach that has been successfully applied to identify unknown ligand binding sites [47]. Our results showed a significantly larger and more varied accumulation of probes within the transmembrane region, with a main site located at the extracellular interface of NisE with NisG, two sites located at the lateral clefts of the transporter, and one minor peripheral site on NisG. Based on the size of the sites, it is likely that the minor site II is not relevant for nisin interaction, while the other sites (Site I, III, and IV) are more likely to provide a marked interaction surface for a peptide the size of nisin.

The results derived from co-solvent MD simulations paint a broader and clearer picture when seen together with the structural prediction of the binding mode of C-Nis. In the latter, all the runs resulted in the peptide being placed on the extracellular face of the interface between NisE and NisG, perfectly matching site I, which was mostly defined by its preference for the backbone mimicking probes (acetone and N-acetamide), supporting the idea that this could indeed be a functionally relevant site, optimized to interact with linear peptides. On this site, the two possible N-to-C orientations along the cleft were observed, but one was 2.8-fold more frequently sampled. This opens the possibility that the rest of the molecule rests in one of the lateral clefts, which have been speculated to be relevant in other peptide-transporting ABC transporters [31], and would agree with either of the lateral sites observed in the co-solvent simulations. The most populated configuration of C-Nis leaves the N-terminus, and therefore the rest of the nisin, in a position where it would be possible to interact with site III. Although this site is smaller than site IV, a small density of acetone and benzene is present, possibly bridging site III to the larger site I, allowing it to form a cohesive binding interface spanning from the top of the transporter to its lateral cleft.

This putative binding interface makes sense in light of the experimental data regarding how both nisin and NisFEG work. Structural studies of the mechanism of action of nisin have shown that the lanthionine rings A, B, and C form a cage surrounding the target molecule, lipid II, whereas the C-terminus of the molecule remains flexible, connected by a hinge region, and is later involved in the formation of bacteriolytic pores [48]. Mutational analysis of nisin variants has shown that both the last six amino acids and the last lanthionine rings D/E are required for NisFEG to effectively protect the cells [26], implying that there must be a direct association between the transporters and these regions of nisin. Since these elements are not bound to lipid II and remain highly flexible, there is a higher likelihood of encountering and binding to NisFEG, thus initiating the first step for extrusion from the membrane. Therefore, these two essential regions, the C-terminus and the last rings, might bind to the identified Sites I and III, respectively.

A more detailed analysis of the putative binding modes of C-Nis reveals that, even though the peptide sits at the middle of both NisG and NisE, the interaction is asymmetric, with more residues from NisE in close contact to the peptide. His31 of the C-terminus is surrounded by aromatic residues from NisE, a result that is consistent across the three largest binding mode clusters. As previously mentioned, all of the binding modes assessed would position the rest of nisin in a position to make interactions with site III, which is also located solely in NisE. This reinforces the asymmetry of function observed for the transmembrane chains. On the one hand, only NisG showed the capacity of forming interactions with the E-loop, on the other hand, NisE seems to dominate the putative interactions with nisin, suggesting that there might be a specialization of roles for each transmembrane chain.. This functional asymmetry could provide an interesting explanation of why there is such a low sequence identity between NisG and NisE; evolution has optimized each chain to perform a specific part of the nisin removal process. Heteromeric complexes formed by homologous chains with low identity are relatively common. For instance, heterodimeric and heterotretrameric ABC transporters are often found in eukaryotic cells, where subunit heterogeneity has been proposed as a means for regulating and fine-tuning the function of the complex [49, 50]. Because the efficacy of a lantibiotic is inherently constrained by the capacity of the producing cell to withstand the toxicity of its own product, there is a strong purging, selective pressure driving the optimization of the immunity-related proteins. Therefore, fine-tuning each transmembrane subunit to drive a specific part of the process could be a favorable evolutionary adaptation.

It is important to note that the large surface and hydrophobic nature of the putative contacts between NisE and NisG make it difficult to experimentally probe the sites, since probably the binding is driven by the sum of the interactions rather than dominated by single residues. Thus, mutational assays might not give conclusive data, and any effect on transporter activity might be confounded with other issues, such as stability and membrane translocation.

Taken together, our results represent the first step in constructing a structural hypothesis of how the LanFEG family of ABC transporters, and more specifically NisFEG, confer immunity to lantibiotic-producing bacterial strains. This mechanism of action is summarized in Figure 8. Nisin, initially bound to lipid II, has its C-terminus free and exposed, which can be recognized by the sites present on NisE. Then, ATP hydrolysis powers conformational changes that are transmitted through the E-loop – NisG association toward the rest of the transporter and enable the removal of nisin from the membrane.

**Figure 8.**
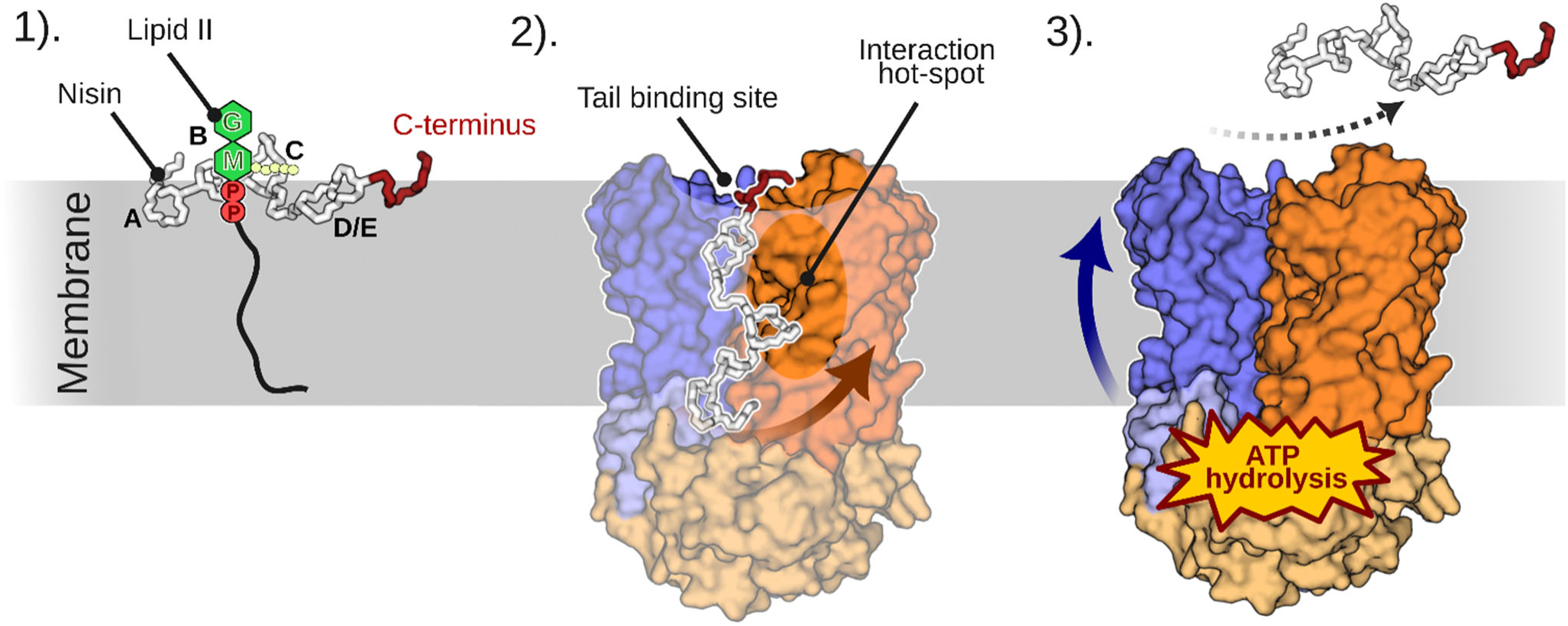
Scheme of a hypothetical mechanism of action for NisFEG. 1) Nisin is bound to Lipid II, but the formation of larger oligomers that form bacteriolytic pores has not yet taken place, therefore, the D/E rings and the C-terminus of nisin move freely. 2) Nisin is first recognized by NisFEG via an interaction of the C-terminus of nisin with the binding sites on NisE (orange arrow). 3) An ATP-powered conformational rearrangement orchestrated by contacts between the E-loop and NisG drives the extrusion of nisin to the extracellular medium (blue arrow), freeing Lipid II and preventing the formation of pores.

## Materials and Methods

### Modeling of NisFEG

A structural model of the transmembrane region involving the protein chains NisE (NCBI: WP_014570414.1) and NisG (NCBI: WP_017864242.1) and NisF in the presence of ATP-Mg was generated using AlphaFold3. The quality of the models was assessed by the metrics of pLDDT and PAE, and by stereochemical analysis with a Ramachandran plot (generated using: https://github.com/Joseph-Ellaway/Ramachandran_Plotter).

The search for further homologous proteins was performed using the FoldSeek [31] webserver with default parameters and the best resulting NisE and NisG models generated by AlphaFold3 as input. The structure of the PmtCD transporter (PDB ID 6XJH) was used as a template for further modeling of the whole complex.

### Conventional and cosolvent all-atom molecular dynamics simulations

The protonation state of the protein was assigned using Propka in Schrödinger [61]. The model was embedded in a DMPC membrane bilayer using PACKMOL-Memgen [62] and TIP3P as the water model. The orientation of the protein within the membrane was previously determined using the OPM-PPM webserver [58]. Topology and coordinates files were generated using tleap [63]. The forcefield ff19SB [64] was used for the protein and lipid17 [65] for the membrane. The parameters for ATP-Mg were taken from the Bryce database [66, 67]. Counterions were added to maintain the electroneutrality of the system. All simulation work was performed using Amber22 [68]. The total system contained 139,261 atoms, made out 31,616 water molecules. 315 lipids, and 33 ions. The initial minimization of the system was performed by adapting the protocol from Roe & Brooks [69], consisting of a total of twelve individual stages. First, 5000 steps of steepest descent minimization followed by 15000 steps of conjugated gradient minimization with positional restraints with a force constant of 5 kcal mol^-1^ Å^-2^ placed on all solute atoms was carried out. Then, the system was heated from 0 to 300 K in an NVT ensemble in a window of 20 ps with a time step of 1 fs, using a Langevin thermostat [70] with a gamma coefficient of 1 ps^-1^, keeping the same restraints in place. Then three successive minimizations of 1000 steps of steepest descent were performed, reducing the restraints on the solute atoms to 2, 0.1, and 0 kcal mol^-1^ Å^-2^. Afterward, an NPT run of 300 ps with a time step of 1 fs was performed, using an anisotropic Monte-Carlo barostat with a pressure relaxation time of 2 ps and a constant temperature of 300 K with a restraint of 1 kcal mol^-1^ Å^-2^ over solute atoms, followed by another identical run but with a lower restraint of 0.8 kcal mol^-1^ Å^-2^. Five subsequent NPT runs of the same length and time steps were performed, placing the restraints only on the protein atoms and decreasing the force constant by 0.2 kcal mol^-1^ Å^-2^ in each run until reaching 0 kcal mol^-1^ Å^-2^. Production runs were performed for 1 µs with a time step of 2 fs using the GPU-accelerated implementation of pmemd [71]. The first 10 ns of production were run with a positional restraint of 0.1 kcal mol^-1^ Å^-2^ on protein atoms and a width of the non-bonded skin of 5 Å and discarded from further analysis. Five independent replicas were carried out saving the coordinates of the system every 100 ps. All simulations were performed using the SHAKE algorithm [72] to constrain the length of bonds to hydrogens and a non-bonded cut-off of 10 Å.

Distances between residues throughout the simulations were calculated using the *distance* command in cpptraj [73]. The distances were calculated between the centre of mass of the carboxylic oxygens of Glu76 and the centre of mass of nitrogen and oxygen of the amide group from either Asn72 or Gln75, or the sidechain nitrogen from Lys129. A per-residue cross-correlation matrix was calculated using the *matrix* command in cpptraj. To suppress spurious correlations, we used only the 80% of the total frames with the lowest RMSD with respect to the starting configuration,

For cosolvent MD simulations, the systems were prepared using acetone, benzene, methanol, or *N*-methylacetamide as a probe. The placement of the probe molecules was performed using PACKMOL-Memgen in a ratio of 1 molecule of co-solvent per 99 molecules of water, using the same membrane and solvent as described previously. The parameters for each co-solvent are included within PACKMOL-Memgen [62]. Five independently packed systems were generated for each probe to randomize the initial position of the probes. The same minimization, thermalization, and pressurization protocol as previously described was used. Five independent production replicas per random packing were carried out, utilizing the same conditions as previously described, but with a production run length of 500 ns. The density distributions for each probe were calculated by first aligning the backbone atoms of the protein using the *rms* command in cpptraj and wrapping the solvent and cosolvent molecules with the *autoimage* function. Densities were calculated using the *volmap* tool of VMD [74] with a resolution of 1 Å and an atom size of 3 Å. To assess the convergence of the cosolvent sampling, the density calculation was repeated at different total simulation times to ensure the invariance of the density distribution. Also, the radial distribution function for each probe was calculated against the Cα atoms of the protein in a window of 100 Å, using the *radial* command on cpptraj [73], ignoring intramolecular interactions.

### Modeling of NisFEG bound to C-Nis

Models of the full NisFEG complex in the presence of ATP-Mg and the last six amino acids of nisin were generated using Boltz-2 [76]. 600 independent runs were performed, with 20 recycle steps. Data processing was carried out using SciPy in Python.

### Site-directed mutagenesis, protein expression, and measurement of NisFEG activity

NisE, NisG, and NisF (or the inactive variant NisF_H181A_, to generate a sensitive control) were cloned into a pILSV shuttle vector as previously described elsewhere [26]. Site-directed mutagenesis was performed by PCR using Phusion DNA polymerase, the template NisFEG-pILSV vector, and a mutagenic primer pair (Table 2). The mutated plasmids were transformed into *L. lactis* NZ9000 by electroporation at 1 kV, 25 μF, 5.0 msec.

**Table 2.**
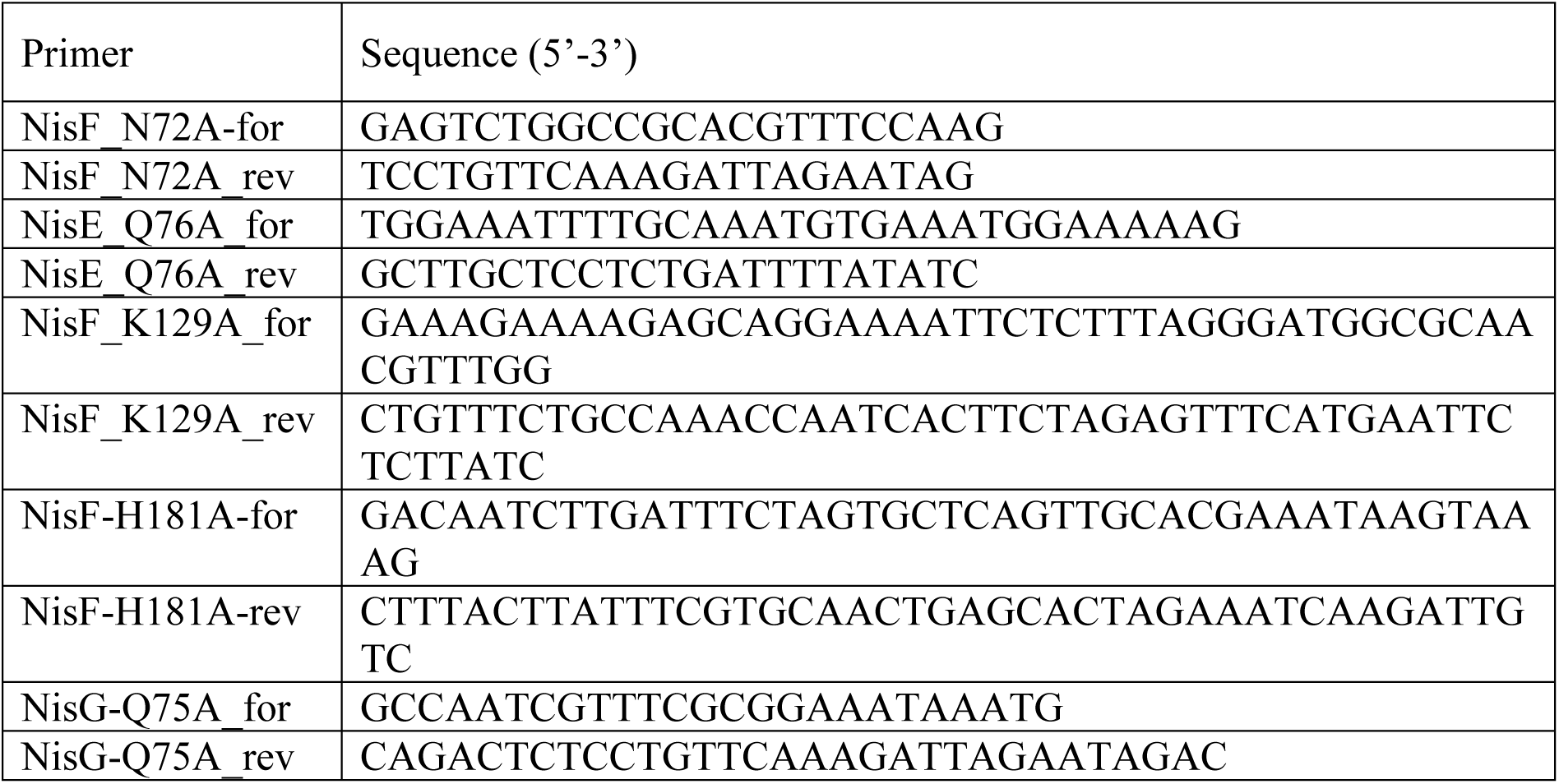
Mutagenic primers used in the present study.

Nisin was purified and quantified as described elsewhere [77]. The transformed *L. lactis* strains harbouring either the WT or a variant were grown overnight in GM17 media supplemented with 10 µg / mL chloramphenicol in the presence of 1 ng / mL of nisin for inducing expression. The next day, the cells were diluted to an OD_600_ of 0.1 in fresh GM17 media and incubated for 30 minutes at 30 °C. Then, 150 µL of bacterial sample were loaded into a 96-well plate alongside 50 µL of nisin solution. The samples were incubated for 5-7 hours at 30°C, and the OD_600_ was measured. The activities of NisFEG, control strain, and mutants were calculated as described in previous work [71].

## ASSOCIATED CONTENT

### Supporting Information

Supporting Information is available free of charge at: [XXX: Add link to paper.]

Additional simulation data shown in supplementary figures and supplementary references (PDF).

### Author Contributions

Conceptualization, PAC, HG; methodology, PAC, SS-V (modeling and simulations) & JG, CM (biochemistry); writing - original draft preparation, PAC, HG; writing - review and editing, JG, CM, SHJS; data analysis and interpretation, PAC, JG, SHJS, HG; supervision and project administration, SHJS, HG; funding acquisition, SHJS, HG. All authors have read and agreed to the published version of the manuscript.

### Funding Sources

This research was funded by grants from the Deutsche Forschungsgemeinschaft [DFG, German Research Foundation; 270650915/GRK 2158 (to HG, SHJS)]. The Center for Structural Studies is funded by the Deutsche Forschungsgemeinschaft [DFG; 417919780 (to SHJS)] and is part of StrukturaLink Rhein-Ruhr, which is funded by the Deutsche Forschungsgemeinschaft [DFG; 573727698].

### Notes

The authors declare no competing financial interest.

## Acknowledgements

We are grateful for computational support and infrastructure provided by the “Zentrum für Informations- und Medientechnologie” (ZIM) at the Heinrich Heine University Düsseldorf and the computing time provided by the John von Neumann Institute for Computing (NIC) to HG on the supercomputer JUWELS at Jülich Supercomputing Centre (JSC) (user ID: VSK33). We thank Martina Wesemann from the Institute of Biochemistry for excellent support.

## DATA AND SOFTWARE AVAILABILITY STATEMENT

The software used for generating the models is in the public domain. The Schrödinger software is available here: https://www.schrodinger.com/. The AMBER package of molecular simulation codes is available here: https://ambermd.org/. Source data, input files, and processing scripts are available at https://researchdata.hhu.de.

## Notes

### Competing Interest Statement

The authors have declared no competing interest.

### Summary of Updates

The paper now presents results only derived from models generated by AlphaFold3, as these show better scores and also show significant structural differences to our original models. All of the mutational data have been reinterpreted in light of the information inferred from the new models. All interactions between residues were quantified in terms of hydrogen bonds. A comparison with a model in the apo state was included in the Supporting Information. The Supporting Information was expanded to include additional figures. A new section dealing with the prediction of the binding mode of the terminal segment of nisin was incorporated.

