## Supporting Information for "Mechanistic Insights into the Structural Asymmetry of the LanFEG Transporter NisFEG in Lantibiotic Immunity"

### Supplementary Figures

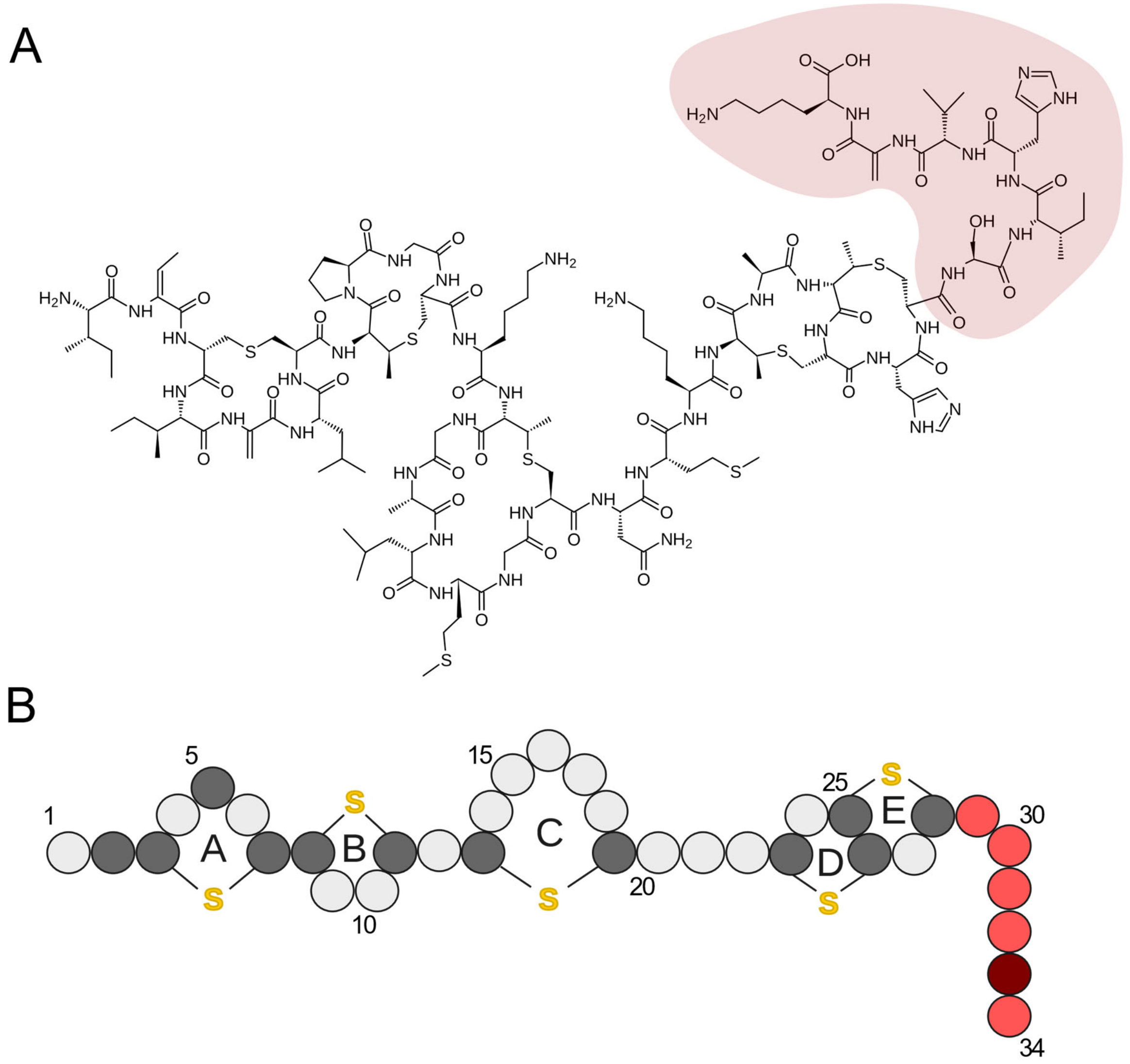


**Supplementary Figure 1. Structure of Nisin.** A) Chemical structure of Nisin. The last six residues, shown to be essential for NisFEG-conferred immunity, are highlighted with a red shade. B) Schematic representation of Nisin. Each bead represents an individual amino acid. Post-translationally modified amino acids are highlighted in a darker shade. Thioether bonds are shown as yellow letters. Each lanthionin ring is labeled with a letter from A to E. The last six residues are shown in red, as in A).

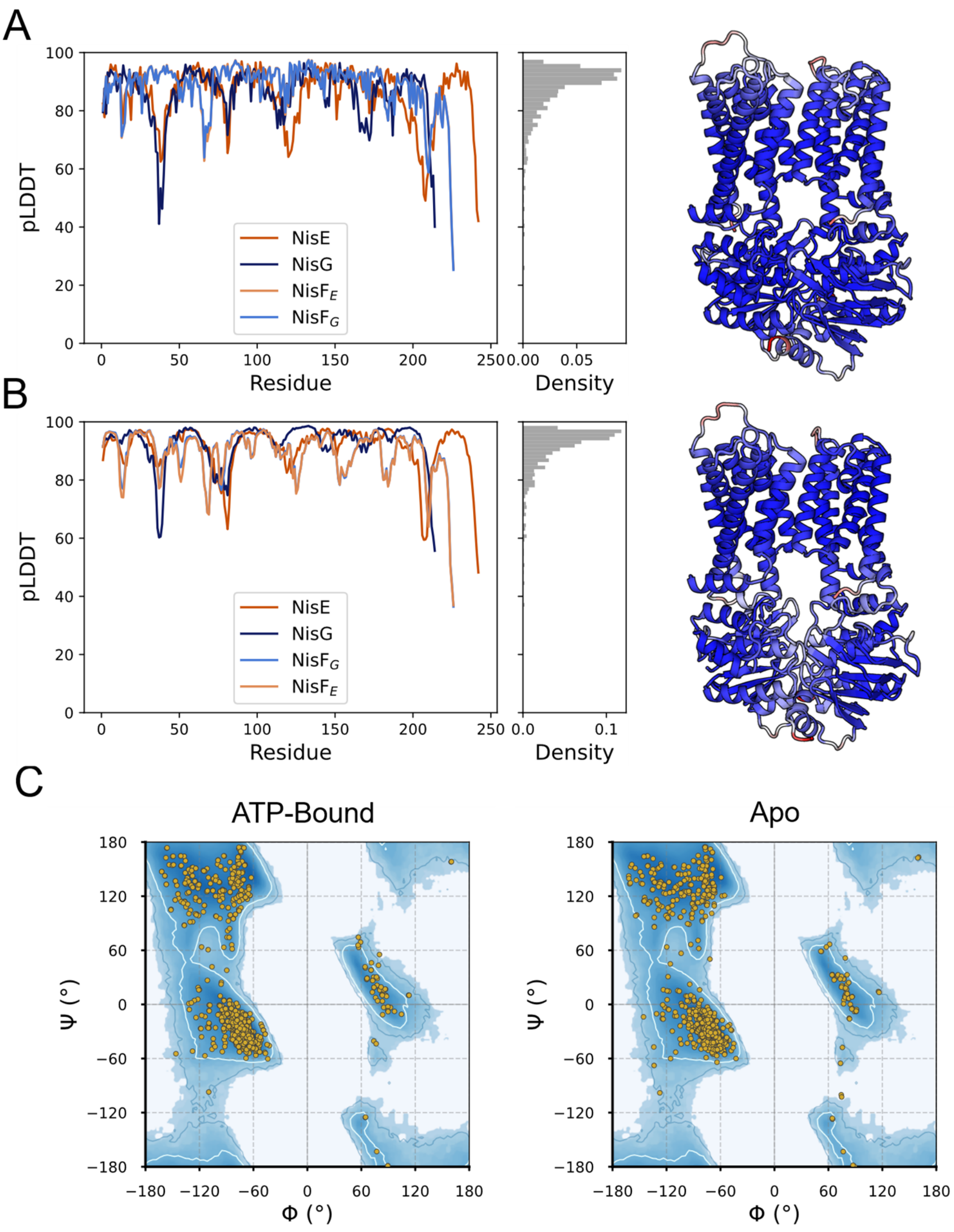


**Supplementary Figure 2: Model quality assessment of NisFEG models in their apo and ATP-bound states.** A) The left panel shows the pLDDT per residue for each one of the individual chains of the ATP-bound model, accompanied by a histogram showing the overall score distribution; the right panel shows the same data, mapped onto the transporter structure. B) Same as in A), but for the apo model. C) Ramachandran plot for each model.

26

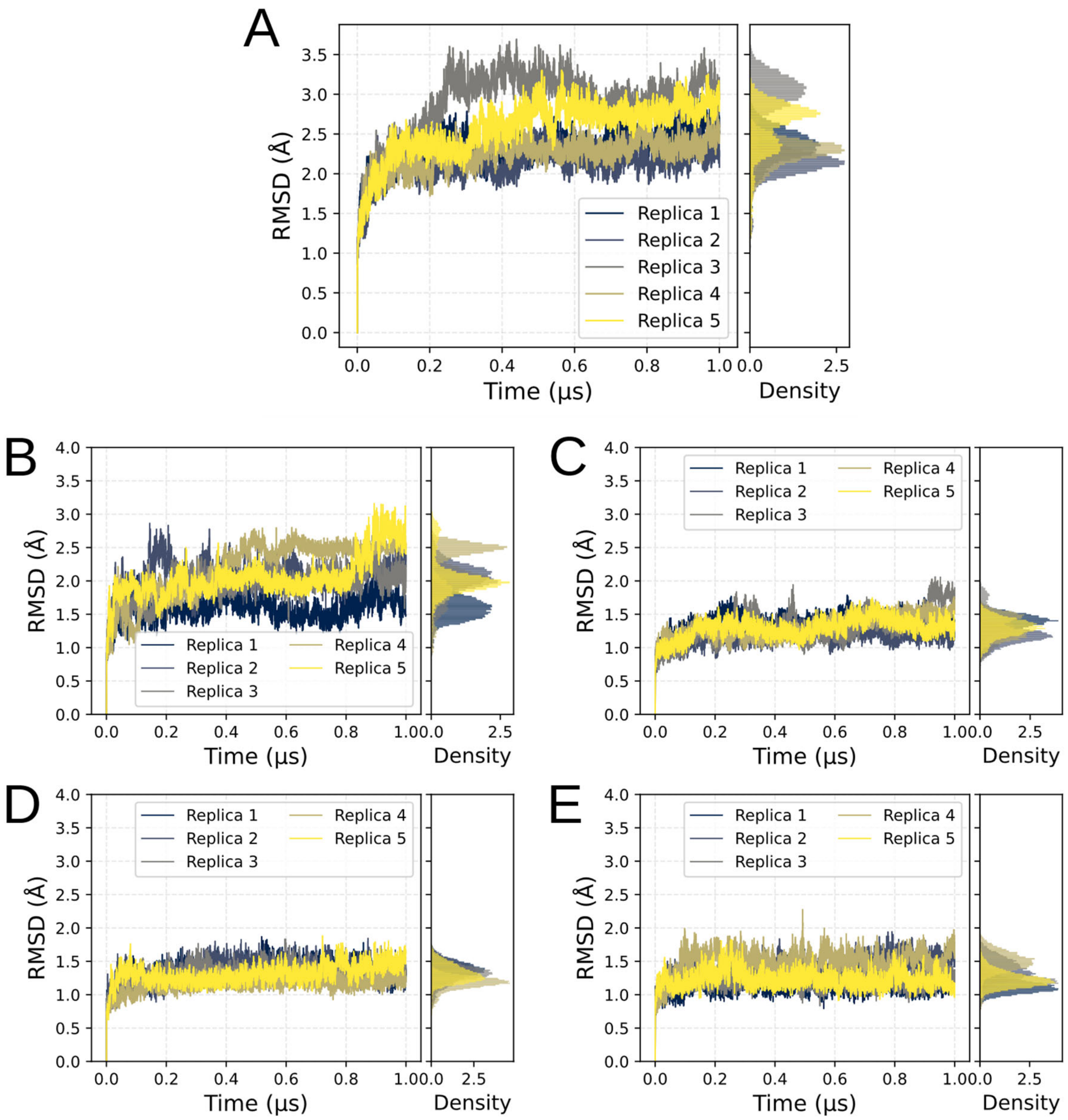


**Supplementary Figure 3. RMSD of backbone atoms of NisFEG with ATP-Mg over five MD simulation replicas.** Each plot shows the RMSD of the backbone atoms versus the simulation time in the left, and an aggregate histogram at the right. A) RMSD for the full NisFEG complex. B) RMSD for the NisE chain. C) RMSD for the NisG chain. D) RMSD for the $\text{NisF}_{\text{E}}$ chain. E) RMSD for the $\text{NisF}_{\text{G}}$ chain.

27

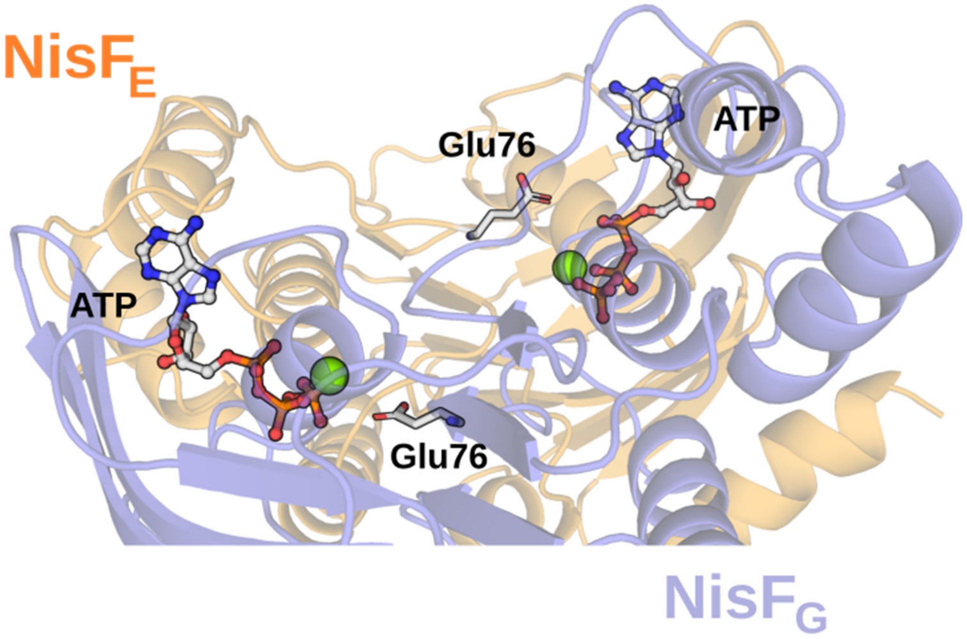


**Supplementary Figure 4. Residue Glu76 of the central E-loop does not interact with Mg-ATP.** Neither of the Glu76 residues (shown as white sticks) makes direct interactions with the Mg-ATP ligands in the NBDs (shown as green spheres and grey sticks, respectively).

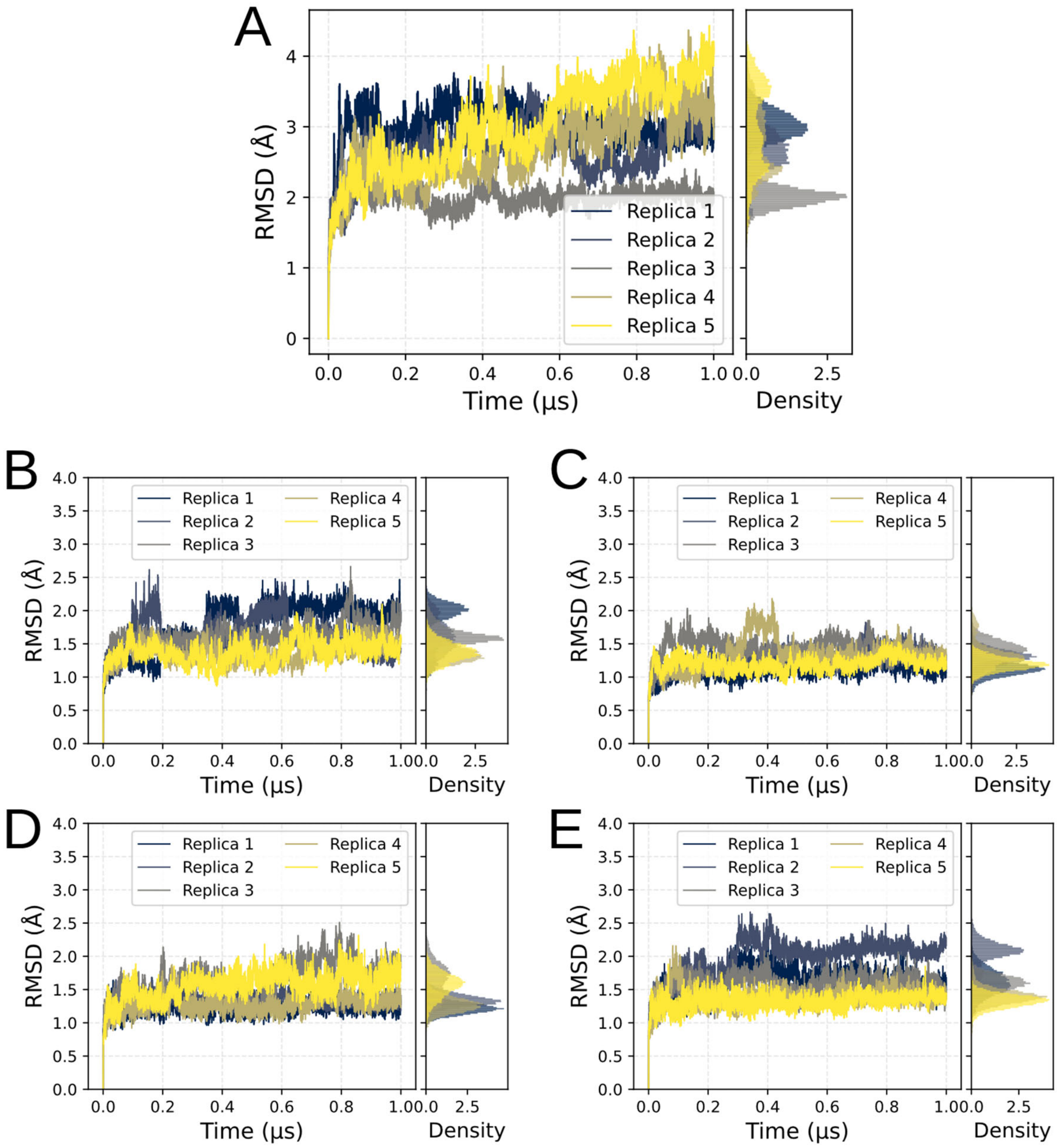


**Supplementary Figure 5. RMSD of backbone atoms of NisFEG in the apo state over five MD simulation replicas.** Each plot shows the RMSD of the backbone atoms versus the simulation time (left), and an aggregate histogram (right). A) RMSD for the full NisFEG complex. B) RMSD for the NisE chain. C) RMSD for the NisG chain. D) RMSD for the $NisF_E$ chain. E) RMSD for the $NisF_G$ chain.

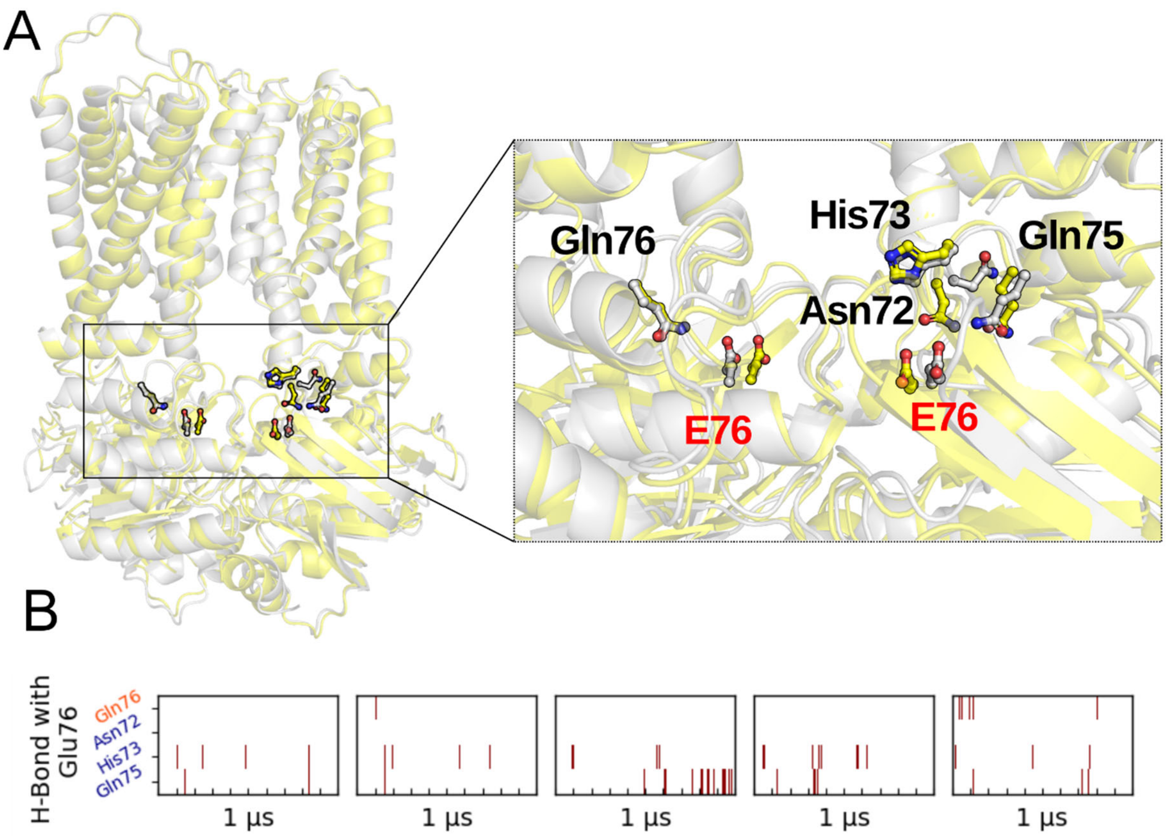


**Supplementary Figure 6. The absence of ATP does not drastically affect the behavior of the E-loop.** A) Structural alignment of the ATP-bound model (yellow) with the apo-state model (white); the ATP molecules were hidden for clarity. The left panel shows a general view of the transporter, while the right panel shows a zoom into the E-loop E76. B) Hydrogen bond pattern for residues. Time-series showing the presence of hydrogen bonds between Glu76 of the E-loop in NisF$_G$ (marked in blue on the y-axis) and NisF$_E$ (marked in orange on the y-axis). The red stripes represent when the residue pairs are forming an effective hydrogen bond (cutoff: 3.5 Å between donor and acceptor and 135° for the acceptor-hydrogen-donor angle) across each replica.

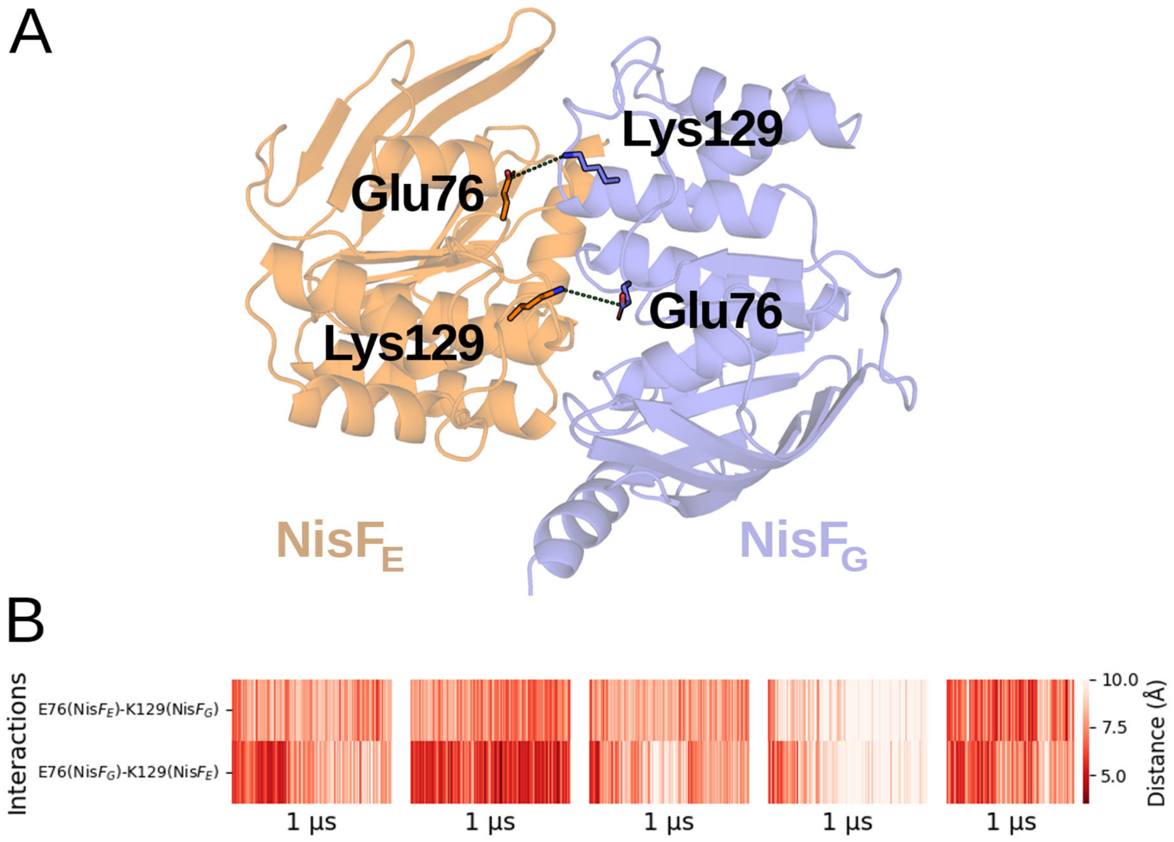


**Supplementary Figure 7. The conserved E76 also provides stabilization for the NBD assembly.** A) Interactions between K129 and E76 in the $NisF_E$ (orange) and $NisF_G$ (blue) subunits. B) Time series of the distance between each K129 - E76 pair for each replica.

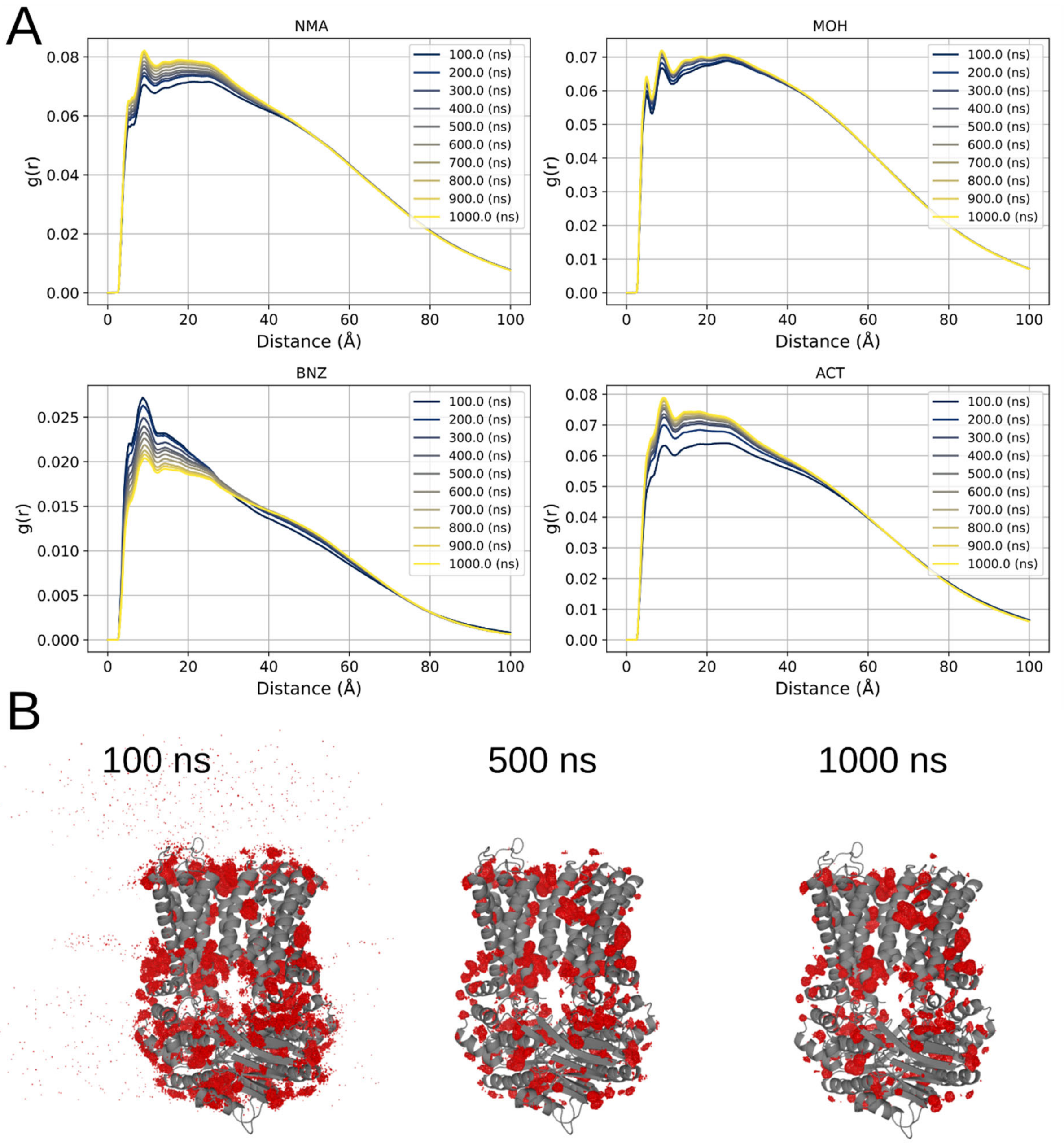


**Supplementary Figure 8. Assessing the convergence of cosolvent MD simulations.** A) Radial distribution function for each one of the probes at different simulation times (NMA: *N*-methylacetamide, MOH: Methanol, BNZ: benzene, ACT: acetone). B) Cosolvent density accumulations over time for a representative probe (methanol) at equal isovalues.
